# Thymosin β4 preserves vascular smooth muscle phenotype in atherosclerosis via regulation of Low Density Lipoprotein Related Protein 1 (LRP1)

**DOI:** 10.1101/2021.11.30.470548

**Authors:** Sonali Munshaw, Andia Redpath, Benjamin Pike, Nicola Smart

## Abstract

**Objective:** Atherosclerosis is a progressive, degenerative vascular disease and a leading cause of morbidity and mortality. In response to endothelial damage, platelet derived growth factor (PDGF)-BB induced phenotypic modulation of medial smooth muscle cells (VSMCs) promotes atherosclerotic lesion formation and destabilisation of the vessel wall. VSMC sensitivity to PDGF-BB is determined by endocytosis of Low density lipoprotein receptor related protein 1 (LRP1)-PDGFRβ complexes to balance receptor recycling with lysosomal degradation. Consequently, LRP1 is implicated in various arterial diseases. Having identified Tβ4 as a regulator of LRP1-mediated endocytosis to protect against aortic aneurysm, we sought to determine whether Tβ4 may additionally function to protect against atherosclerosis, by regulating LRP1-mediated growth factor signalling.

**Approach and Results:** By single cell transcriptomic analysis, *Tmsb4x*, encoding Tβ4, strongly correlated with contractile gene expression and was significantly down-regulated in cells that adopted a modulated phenotype in atherosclerosis. We assessed susceptibility to atherosclerosis of global Tβ4 knockout mice using the ApoE^-/-^ hypercholesterolaemia model. Inflammation, elastin integrity, VSMC phenotype and signalling were analysed in the aortic root and descending aorta. Tβ4KO; ApoE^-/-^ mice develop larger atherosclerotic plaques than control mice, with medial layer degeneration characterised by accelerated VSMC phenotypic modulation. Defects in Tβ4KO; ApoE^-/-^ mice phenocopied those in VSMC-specific LRP1 nulls and, moreover, were underpinned by hyperactivated LRP1-PDGFRβ signalling.

**Conclusions:** We identify an atheroprotective role for endogenous Tβ4 in maintaining differentiated VSMC phenotype via LRP1-mediated PDGFRβ signalling.

## Introduction

Atherosclerosis is a dynamic arterial disease, initiated by endothelial dysfunction^1^. Subendothelial lipid retention provokes a maladaptive inflammatory response and leads to the formation of atheromatous plaques, characterised by a necrotic core and enveloped by a fibrous cap^2^. The resultant loss of vessel wall integrity is detrimental, predisposing affected individuals to an array of complications, including aneurysm and thrombosis. Whilst gradual occlusion of vessels may precipitate as claudication, the principal clinical consequences arise from thrombotic and embolic events associated with unstable plaques that are prone to erosion or rupture^3^. The consequent cardiovascular morbidities – which include coronary heart disease, ischaemic stroke and peripheral artery disease – remain a major societal burden, for which interventions beyond lifestyle change are relatively limited^4^.

Plaque stability is determined by opposing degenerative and regenerative processes. Degradation of the medial layer is mediated by the proteolytic actions of macrophages and potentiated by pro-inflammatory signalling, while the reparative response comprises vascular wall remodelling and synthesis of extracellular matrix (ECM), processes ascribed to vascular smooth muscle cells (VSMCs)^5^. In healthy tissue, VSMCs are chiefly localised to the tunica media, maintaining a fully differentiated contractile phenotype to regulate vascular tone. This state is defined by expression of proteins of the contractile apparatus, including smooth muscle myosin heavy chain (*Myh11*), alpha-smooth muscle actin (*Acta2*) and transgelin (*Tagln*)^6^. However, in atherosclerosis, environmental triggers initiate a growth factor-mediated phenotypic transition causing VSMCs to de-differentiate, down-regulate contractile marker expression, proliferate, and migrate to the intimal layer^7^. Modulated, so-called ‘synthetic’ VSMCs, are proposed to adopt a phenotype that is either macrophage-like, pro-inflammatory, and destabilising, or fibroblast-like, synthetic, and stabilising^8^. Although reparative in the short term, chronic VSMC dedifferentiation exacerbates inflammation to promote atherosclerotic lesion development, leading to arterial stiffness and stenosis^9^. Therefore, in the main, inhibiting VSMC phenotypic transformation has been shown to attenuate progression of vascular disease^10^.

Various signalling pathways, including transforming growth factor (TGF)β, NOTCH, Wnt and Sonic hedgehog (Shh), promote mural cell differentiation during development and serve to maintain VSMC contractile state in the adult vasculature (summarised in ^11^). Conversely, the contractile-synthetic phenotypic modulation of VSMCs is induced primarily by platelet-derived growth factor (PDGF)-BB, levels of which increase markedly in response to endothelial damage^12^. The response of VSMCs to PDGF-BB is determined by the extent of PDGF receptor β (PDGFRβ) exposure at the cell surface, which is controlled by endocytic trafficking in complex with the co-receptor, Low density lipoprotein (LDL) receptor related protein 1 (LRP1)^13^. Following internalisation and transduction of signals to nuclear effectors, LRP1-PDGFRβ complexes may be recycled to the cell membrane or targeted for lysosomal degradation, to potentiate or attenuate downstream signalling activity, respectively^14^. We recently defined a key role for the small peptide Thymosin β4 (Tβ4) in regulating LRP1-PDGFRβ trafficking to control VSMC phenotype and protect against abdominal aortic aneurysm (AAA)^15^. This study demonstrated a postnatal VSMC-autonomous role for Tβ4 to maintain VSMC differentiation for vascular homeostasis beyond its developmental role to induce aortic VSMC differentiation^16^. Because of its central role in controlling the sensitivity of VSMCs to growth factors, LRP1 has been implicated through genome-wide association and animal studies in protection against a range of arterial diseases, including AAA^17^, carotid^18^ and coronary artery disease^19^. We therefore sought to determine whether Tβ4 functions to protect against atherosclerosis by regulating LRP1-PDGFRβ signalling to preserve VSMC phenotype. In a murine model of atherosclerosis, Tβ4-null mice displayed an exacerbated phenotype, relative to controls, forming large plaques in the aortic sinus and descending aorta. Notably, accelerated disease progression closely phenocopies that observed in VSMC-specific *Lrp1* knockout mice. While inflammation in the aorta was unaltered in Tβ4 mutants, augmented plaque formation was characterised by accelerated VSMC phenotypic modulation and elastolysis, underpinned by dysregulated LRP1-PDGFRβ signalling. These findings support a common mechanism through which the Tβ4-LRP1 axis controls growth factor signalling, VSMC differentiation and protection against aortic diseases.

## Methods

### Mouse lines

Mice were housed and maintained in a controlled environment and all procedures involving the use and care of animals were performed in accordance with the Animals (Scientific Procedures) Act 1986 (Home Office, United Kingdom) and approved by the University of Oxford Animal Welfare and Ethical Review Boards. All mice used for the study were maintained on the C57BI6/J background for at least 20 generations. The global Tβ4 Knockout strain, in which exon 2 of *Tmsb4x* locus is deleted, were a kind gift of Martin Turner, Babraham Institute and have been previously described^16^. VSMC-specific Lrp1 knockouts were generated by crossing the previously described floxed Lrp1 line^20^ (obtained from Jax) to homozygosity with Myh11^CreERT221^, to generate Myh11^CreERT2^; Lrp1^fl/fl^. These lines were crossed to homozygosity with ApoE^-/-^mice^22^ (from Jax). Conditional knockout of *Lrp1* was induced by oral gavage of 3 week old mice with 3 doses of tamoxifen (80mg/kg) on alternate days.

### Atherosclerosis model

Age-matched (8-12 week old) mice, on the ApoE^-/-^ background, were fed on Western diet (21.4% fat; 0.2% cholesterol; Special Diets Services, Essex, UK) for up to 12 weeks. Mice were weighed weekly and, at harvest, serum collected for cholesterol analysis (by Charles River Laboratories). EdU(5-ethynyl-2’-deoxyuridine), prepared in 0.9% saline, was administered at a dose of 150mg/kg by intraperitoneal injection (4 doses over the 2 weeks prior to harvest).

### Single cell RNA sequencing analysis

10X Chromium data were downloaded as cell–gene count matrices from GSE131776^23^. The scRNA-seq dataset, specifically the tdTomato positive VSMC lineage, was analysed using Seurat^24, 25^ in R, as follows. Cells with more than 8% mitochondrial reads, more than 3500 or fewer than 1000 genes per cell were filtered out. The principal component analysis was used to cluster cells, which were visualized with the UMAP method^26^ and biological samples – Baseline, 8 and 16 week HFD – were integrated using Harmony^27^. Differential expression (DE) analysis Wilcoxon rank-sum test was carried out between biological samples and clusters, to identify upregulated and downregulated genes (Log2FC) with significant adjusted p-value (Bonferroni correction).

### Histological Sample Preparation

For histological and immunohistochemical analyses, mouse aortic sinus, aortic arch and descending aorta were harvested, washed in phosphate-buffered saline (PBS; pH7.4) and fixed in 4% paraformaldehyde (PFA) at room temperature, either for two hours (cryosectioning) or overnight (paraffin embedding). For histological analysis, samples were dehydrated using a series of graded ethanol concentrations (50-100%, each for at least 2 hours), and cleared in butanol overnight. Samples were incubated at 60°C in 1:1 butanol: molten pastillated fibrowax for 30 minutes, then three times in 100% molten wax, each for 4 hours to overnight, before embedding and sectioning (10μm transverse sections). Paraffin sections were stained with Oil red O solution, Hematoxylin and Eosin, Elastic stain kit (Verhoeff van Gieson) and Masson’s Trichrome kit (all from *Sigma-Aldrich*), each according to the manufacturer’s protocol. After imaging, morphology was assessed on sections which had been anonymised and blinded for genotype. Elastin breaks per section were counted manually. Plaque area and aortic diameter were measured using ImageJ. Elastin breaks/score, aortic diameter, TA/AA plaque size are all expressed as the mean of 6 sections per aorta) For immunohistochemical analyses, fixed tissues were rinsed in PBS and equilibrated in 30% sucrose (in PBS) overnight at 4°C. Samples were placed in 50:50 (30% sucrose: Tissue-Tek OCT), then embedded in OCT and chilled at −80°C.

### Immunofluorescence Staining on Cryosections

Frozen sections of descending aorta or aortic sinus (base of the heart) were cut at a thickness of 10 µm, air-dried for 5 minutes and rinsed in PBS. Sections were permeabilized in 0.5% Triton-X100/PBS for 10 minutes, rinsed in PBS and blocked for 1-2 hours (1% BSA, 10% goat serum or donkey serum; 0.1% Triton-X100 in PBS). Sections were incubated with primary antibodies (diluted as below) at 4°C overnight. Sections were washed 5-6 times in 0.1% Triton-X100 in PBS (PBST) then incubated in secondary antibody for 1 hour at room temperature. Sections were washed 3 times in PBST and, incubated with 300nmol/uL 4’,6-diamidino-2-phenylindole (DAPI) in PBS for 5 minutes and rinsed a further twice in PBS. Slides were mounted with 50% glycerol:PBS and imaging was conducted on an Olympus FV1000 confocal microscope or a Leica DM6000 fluorescence microscope. Apoptosis was assessed using the Click-iT TUNEL Alexa Fluor 546 Imaging Assay kit and EdU incorporation by Click-iT EdU Alexa Fluor 546 Imaging Assay kit (both from ThermoFisher), according to the manufacturers’ instructions. Antibodies used for Immunofluorescence: FITC-conjugated Mouse anti-αSMA (1:200; Sigma; F3777); Rabbit anti-Vimentin (1:200; Abcam; ab45939); Rat anti-CD68 (1:100; BioRad; MCA1957); Rabbit anti-Caldesmon (1:250 Abcam; ab32330); Rabbit anti-Thymosin β4 (1:100; Immunodiagnostik; A9520); Rabbit anti-phospho-LRP1 (Tyr4507) (1:100; Santa Cruz Biotechnology, sc-33049); Rabbit anti-phospho-PDGFRβ (Tyr1021) (1:100; Abcam, ab134048); Rabbit anti-HTRA1 (1:100; Abcam, ab38611); Rabbit anti-Lumican (1:200; Abcam, ab168348); AlexaFluor-conjugated secondary antibodies raised against Rat or Rabbit IgG (Invitrogen) were used at 1:200.

### Immunoblotting

Aortic biopsies were lysed in ice cold RIPA buffer (Sigma) supplemented with 1mM dithiothreitol, phosphatase and protease inhibitor cocktail (PhosSTOP and cOmplete protease inhibitor cocktail; Roche). Western blots were performed using a standard protocol. Proteins were transferred by semi-dry transfer onto PVDF membrane. After blocking, membranes were probed with primary antibodies overnight. Antibodies used were Rabbit anti-phospho-Akt (S473) (1:500; Cell Signalling Technology, 4058), Rabbit anti-phospho-ERK1/2 (Thr202/Tyr204) (1:1000; Cell Signalling Technology, 9101), Rabbit anti-eIF4E (1:1000; Cell Signalling Technology, 2067), Rabbit anti-β-actin (1:1000; Abcam, ab8227). HRP-conjugated secondary antibodies were from GE Healthcare (rabbit, NA934) and R&D (sheep, HAF016).

### RNA isolation and gene expression profile by quantitative real time (qRT-PCR)

Total RNA was isolated from the aortic arch or descending aorta of adult mice using the RNeasy Fibrous Tissue Mini kit (Qiagen), according to the manufacturer’s instructions. cDNA was prepared using RT kit (Applied Biosystems). qRT-PCR was performed on a ViiA 7 (Life technologies) using fast SYBR Green (Applied Biosystems). Data were normalised to β-actin expression (endogenous control, after selection of optimal control, in terms of stable expression, from a panel of 6 genes). Fold-changes were determined by the 2−^ΔΔ^CT method^28^.*Primer sequences (5’-3’)*: *β-actin* F: GGCTGTATTCCCCTCCATCG; R: CCAGTTGGTAACAATGCCATGT; *sm22α* F: CAACAAGGGTCCATCCTACGG; R: ATCTGGGCGGGCCTACATCA; *Myh11* F: AAGCTGCGGCTAGAGGTCA; R: CCCTCCCTTTGATGGCTGAG; *Caldesmon* F: GCTCCCAAGCCTTCTGACTT; R: CCTTAGTGGGGGAAGTGACC; *Vimentin* F: TCCAGCAGCTTCCTGTAGGT; R: CCCTCACCTGTGAAGTGGAT; *Il1b* F: CAACCAACAAGTGATATTCTCCATG; R: GATCCACACTCTCCAGCTGCA; tnf*α* F: CACGTCGTAGCAAACCACCAAGTGGA; R: TGGGAGTAGACAAGGTACAACCC

### Statistics

Randomisation of animals to treatment or genotype groups was introduced at the time of introduction of modified diet. Thereafter, tissues were processed and analysed by an independent observer, whilst blinded to treatment or genotype. Statistical analyses were performed with GraphPad Prism Software. For the quantitative comparison of two groups, two-tailed unpaired Student’s t-test was used to determine any significant differences, after assessing the requirements for a t-test using a Shapiro-Wilk test for normality and an F-Test to compare variances. Alternatively, a Mann-Whitney non-parametric test was used. For comparison of three groups or more, a one-way ANOVA with Tukey’s post-hoc test was used. For analyses involving two independent variables, a two-way ANOVA with Bonferroni, Holm-Sidak or Dunnett’s post-hoc test was used. Significance is indicated in the figures, as follows: *: p≤0.05; **: p≤0.01; ***: p≤0.001; ****: p≤0.0001.

## Results

### Phenotypic modulation of VSMCs in atherosclerosis is associated with down-regulation of Tmsb4x

To understand how endogenous Tβ4 levels are altered during disease, and whether they may influence contractile phenotype at the level of individual VSMCs, we first re-analysed published single-cell RNA sequencing (scRNA-Seq) data. The study by Wirka and colleagues utilised VSMC (*Tg*^*Myh11-CreERT2*^) lineage traced mice on the ApoE^−/−^ background, to perform transcriptomic analyses at baseline and following 8 or 16 weeks of high-fat diet (HFD) to induce atherosclerosis^23^. While the authors sequenced the full spectrum of cells in the aortic root and ascending aorta using the 10x Chromium platform, we focussed our analysis on the tdTomato-labelled cells determined to be of VSMC origin (Fig. 1A-C; Online Figure I). Across all time points, three discrete populations of contractile VSMCs could be discerned (ContSMC_1-3), based on differential gene expression, as well as a transcriptomically distinct cluster of modulated VSMCs (ModSMC) and the pericyte population identified in the original study (heat map shown in Online Figure IA). The contractile VSMC clusters exhibited enriched expression of canonical contractile markers, including *Myl9, Acta2, Tmp2*, and *Tagln*, and relatively low expression of so-called ‘synthetic’ markers – such as *Dcn, Lum*, and *Col1a1*. While contractile marker expression was comparatively uniform across all three contractile clusters, ContSMC_2 and ContSMC_3 were distinguished by higher expression of injury-response genes, including *Atf3, Sost, and Junb*, compared with ContSMC_1. Further separation of ContSMC_2 and ContSMC_3 related primarily to a higher expression of cell cycle arrest genes, such as *Gadd45b*, in ContSMC_3. The ModSMC cluster was characterised by a relative abundance of the ‘synthetic’ markers, including *Dcn, Lum*, and *Fn1* – alongside a relative reduction of contractile markers, such as *Acta2, Myh11*, and *Cnn1* (Table I). The majority of enriched genes related to cell migration and extracellular matrix (ECM) functions, and also included markers associated with mesenchymal cell (*Sca1*) and macrophage (*Lgals3*) identity, consistent with previous descriptions of the modulated VSMC phenotype^3, 29, 30^.

**Figure 1.**
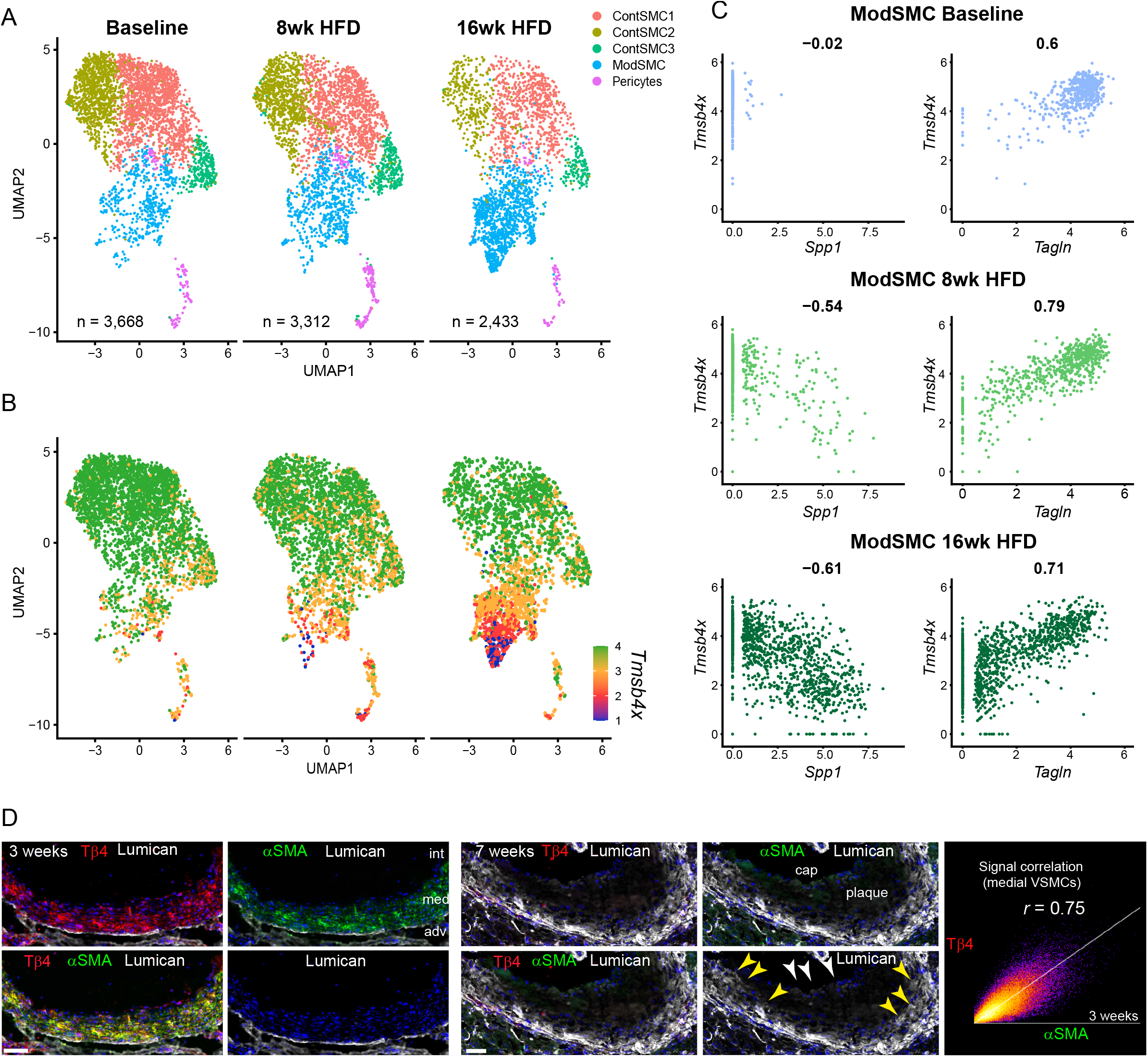
Contractile VSMC Phenotype in atherosclerosis is correlated with expression of *Tmsb4x*. Uniform Manifold Approximation and Projection (UMAP) plots demonstrating transcriptomically distinct contractile (ContSMC1-3) and modulated (ModSMC) VSMC subpopulations of ApoE^-/-^ mice at baseline and after 8- and 16- weeks of high fat diet (HFD; **A**). UMAP showing changes in the expression of *Tmsb4x* across the VSMC subpopulations over the time course of atherosclerosis (**B**). Each dot is an individual cell and gene expression levels are indicated by the colour spectrum, with green representing high, orange medium and red-blue low *Tmsb4x* expression. n=3 for baseline, 8- and 16 weeks. Data re-analysed from ^23^. *Tmsb4x* expression in the ModSMC cluster positively correlated with contractile marker expression, as shown for *Tagln*, and negatively correlated with synthetic markers, exemplified by *Spp1* (**C**). Pearson correlation coefficients are indicated above each plot. Validation by immunofluorescence in aortic sinus sections from ApoE^-/-^ male mice (**D**); Tβ4 and α-smooth muscle actin (α-SMA) abundantly co-expressed in medial layer VSMCs at 3 weeks’ HFD but expression of both declined by 7 weeks’ HFD. Across both time points, Coloc2 quantified Pearson correlation coefficients ranged 0.64-0.75. Lumican was up-regulated with disease progression in medial (yellow arrowheads) and cap VSMCs (white arrowheads), which were largely devoid of Tβ4 and α-SMA. Scale bar in **D**: 50μm.

The contractile-synthetic VSMC phenotypic transition associated with plaque development was evidenced by a shift in cell composition after HFD feeding (Figure 1A and Online Figure IB). From a baseline of 83% ContSMCs and 14.2% ModSMCs, the modulated population expanded, to account for 20.3% and 47.5% of VSMC lineage derivatives at 8 and 16 weeks, respectively, with a corresponding decrease of the contractile fraction to 73.6% and 49.9%, respectively. Even at baseline, levels of *Tmsb4x*, the transcript encoding Tβ4, were marginally lower in the ModSMC cluster, compared with ContMSCs (Figure 1B and Online Figure IC). With HFD, a striking, progressive reduction in *Tmsb4x* expression was observed in this cluster, to the extent that *Tmsb4x* featured among the top 20 down-regulated genes, alongside canonical contractile markers, such as *Tnnt2, Acta2, Myl9, Myh11* and *Myl6* (log2FC<1, baseline vs 16 weeks’ HFD; Table II). With disease progression, *Tmsb4x* expression across the expanding ModSMC cluster became strongly correlated with contractile marker expression, as shown for *Tagln* (Pearson correlation coefficient = 0.79 at 8 weeks; 0.71 at 16 weeks; Figure 1C), with a negative correlation emerging between *Tmsb4x* and synthetic markers, exemplified by *Spp1*, only after HFD feeding (Pearson correlation coefficient = −0.54 at 8 weeks; −0.61 at 16 weeks Figure 1C). This relationship was validated at the protein level in aortic sinus sections from ApoE^-/-^ male mice after a period of HFD. Both Tβ4 and contractile protein α-smooth muscle actin (α-SMA) were abundantly expressed at baseline and early disease stages (3 weeks’ HFD, Figure 1D) and both declined with disease progression (7 weeks’ HFD, Figure 1D). Tβ4 expression strongly correlated with levels of α-SMA in medial layer VSMCs (Pearson correlation coefficient = 0.69 (range 0.64-0.75 across both time points), quantified using Coloc2 in Image J; Figure 1D). Consistent with the scRNA-Seq, ModSMCs increased in number with disease progression and were more prominent within the lesion and were largely found within the fibrous caps. *Lum*, encoding the ECM protein Lumican, emerged from the scRNA-seq as a useful marker of modulated VSMCs (Online Figure IA, C, Table I). At baseline and early disease stages, Lumican expression was largely confined to the external elastic membrane and barely detectable in VSMCs (Figure 1D). However, with disease progression, a progressive up-regulation of Lumican was observed in an increasing number of VSMCs in the medial layer and fibrous cap (Figure 1D). Lumican-expressing modulated VSMCs had substantially lower levels of Tβ4 and αSMA (Figure 1D; shown at 7 weeks’ HFD;). Indeed, a striking inverse relationship between *Tmsb4x* and *Lum* is apparent within the ModSMC cluster across the disease time course (Online Figure IC). These data strongly support a correlative relationship between high Tβ4 levels and maintenance of the contractile VSMC phenotype, consistent with its roles in promoting mural cell differentiation during development^16, 31, 32^, in vascular homeostasis postnatally and in contractile VSMC preservation in a murine AAA model^15^.

### Loss of Tβ4 predisposes to atherosclerosis

To determine whether a reduction in Tβ4 levels, as occurs during atherosclerosis, may contribute towards the modulation of VSMC state, we investigated the phenotype of Tβ4-null male mice (Tβ4^-/Y^) in a model of hypercholesterolaemia, by crossing onto the ApoE^-/-^ background^22^ and feeding a HFD (21.4% fat; 0.2% cholesterol) for up to 12 weeks. To evaluate the hypothesised common function for Tβ4 and LRP1 in regulating VSMC phenotypic modulation, we generated a VSMC-specific *Lrp1* knockout (KO) line (Myh11^Cre^; Lrp1^fl/fl^; ApoE^-/-^; tamoxifen-induced at 3 weeks of age), for comparison with Tβ4 KO mice. Smooth muscle-specific *Lrp1* deletion, whether constitutively or inducibly targeted, strongly predisposes mice to develop atherosclerotic and aneurysmal disease^13, 20, 33^. Advanced atherosclerotic lesions, containing large necrotic cores, were detected in the aortic sinus (Figure 2A, C) and descending thoracic aorta (Figure 2B, D) of all mice. By area, plaques were significantly larger in Tβ4^-/Y^; ApoE^-/-^ mice than in Tβ4^+/Y^; ApoE^-/-^ littermate controls and were comparable in size to those of Myh11^Cre^; Lrp1^fl/fl^; ApoE^-/-^ mice (Figure 2A-D). Increased plaque size did not correlate with an increase in weight gain or in cholesterol levels in Tβ4^-/Y^; ApoE^-/-^ mice, compared with Tβ4^+/Y^; ApoE^-/-^, although mice of both genotypes weighed more than Myh11^Cre^; Lrp1^fl/fl^; ApoE^-/-^, both at baseline and after high fat feeding (Online Figure II). Whilst our study initially examined male mice in HFD cohorts, the unanticipated observation of hindlimb paralysis, a possible indication of aortic thrombosis, aneurysm or rupture, prompted us to investigate a cohort of regular chow-fed 8 month old female Tβ4^-/-^; ApoE^-/-^ and age-matched Tβ4^+/+^; ApoE^-/-^ mice. Even without a cholesterol-rich diet, aortas of Tβ4^-/-^; ApoE^-/-^ mice were aneurysmal (>1.5 fold dilated), compared with Tβ4^+/+^; ApoE^-/-^, and contained large atherosclerotic plaques (Figure 2E-G). In view of the increasingly recognised sexual dimorphism in vascular disease, both in patients and in experimental animals^34^, we compared a further cohort of female mice, this time with HFD over an equivalent time course to that used in male cohorts, and determined a significant increase in aortic arch plaque size in Tβ4^-/-^; ApoE^-/-^, compared with Tβ4^+/+^; ApoE^-/-^ mice (Online Figure III). Interestingly, in female mice, as compared to male mice, increased plaque size did correlate with increased weight gain in Tβ4^-/-^; ApoE^-/-^ mice, compared with Tβ4^+/+^; ApoE^-/-^. Taken together, these studies suggest that atherogenesis is accelerated in Tβ4 null mice, regardless of gender, and closely recapitulates the phenotype associated with loss of LRP1.

**Figure 2.**
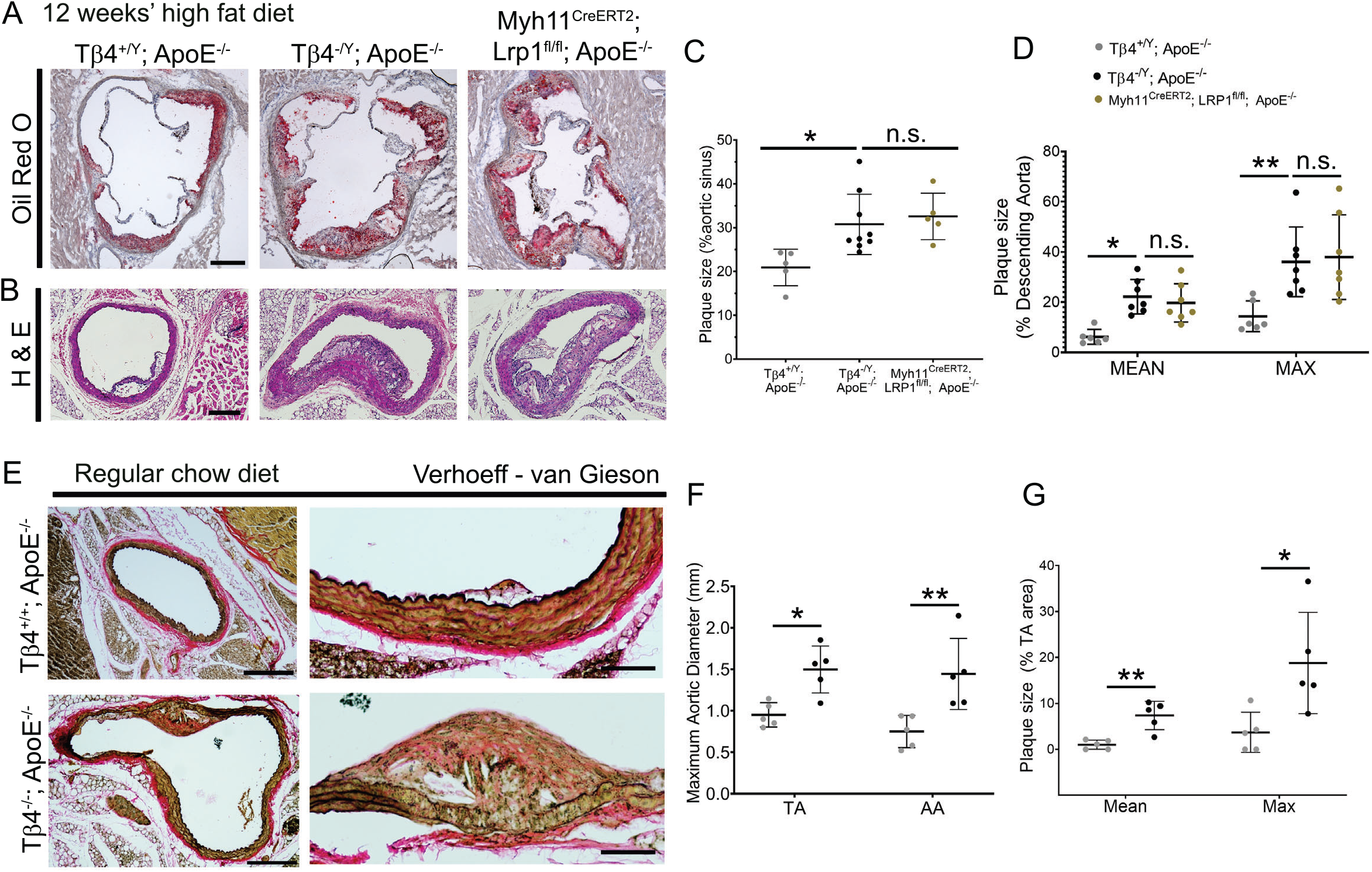
Tβ4-null mice develop larger atherosclerotic lesions. Oil red O staining of aortic sinus plaques from Tβ4^+/Y^; ApoE^-/-^(n=5), Tβ4^-/Y^; ApoE^-/-^ (n=9) and Myh11^Cre^; Lrp1^fl/fl^; ApoE^-/-^ (n=5) mice. (**A**), with quantification of total plaque area in **C**. Plaques in the descending aorta from Tβ4^+/Y^; ApoE^-/-^(n=6), Tβ4^-/Y^; ApoE^-/-^ (n=7) and Myh11^Cre^; Lrp1^fl/fl^; ApoE^-/-^(n=7) mice, assessed by hematoxylin and eosin (H&E) staining, with quantification of total plaque area in **D**. Plaques were measure in 22-week old mice, after 12 weeks of high fat diet. Even without high fat diet, 8 month old females showed aortic dilatation (**E-F**) and larger plaque formation (**E-G**; n = 5). Data are presented as mean ± SD, with each data point representing an individual animal. Significance was calculated using one-way ANOVA with Tukey’s multiple comparison tests (**C, D**) or Mann Witney non-parametric test with Holm-Sidak correction for multiple comparisons (**B, D**). n.s. = not significant; *p ≤ 0.05; **p≤0.01. Scale bars: **A**: 500μm; **B**: 200 μm; **E**: 500μm (magnified panels, right: 150 μm).

### Accelerated plaque formation is not due to exacerbated inflammation from loss of Tβ4

Acknowledging the prominent underlying role of inflammation in the initiation and progression of vascular disease, and the numerous anti-inflammatory roles ascribed to Tβ4^35-37^, we characterised the inflammatory responses of Tβ4^-/Y^; ApoE^-/-^ and Tβ4^+/Y^; ApoE^-/-^ mice during atherosclerosis. Quantitative reverse transcription real-time PCR (qRT-PCR) indicated comparable expression of pro-inflammatory cytokines *Il1b* and *Tnfα* in descending aorta biopsies between genotypes at 3, 7, 9 and 12 weeks’ HFD (Figure 3A). Similarly, levels of pro-inflammatory cytokines, Interferon (IFN)γ, Interleukin (IL)-6 and Tumor necrosis factor (TNF)α, in peripheral blood were not significantly different between genotypes (Figure 3B). This was borne out in the quantification of immune cells recruited to aortic lesions. Monocyte (CD45+CD11b+CD14+) content of the descending aorta was assessed by flow cytometry and found to be comparable between genotypes (Figure 3C). Recruited monocytes were further defined based on expression of the chemokine receptor CCR2, implicated in recruitment of monocyte-derived macrophages to the aorta and development of AAA^38^. The number of recruited CD45+CD11b+CD14+CCR2+ macrophages was indistinguishable between Tβ4^-/Y^; ApoE^-/-^ and Tβ4^+/Y^; ApoE^-/-^ (Figure 3C). Finally, we assessed macrophage/foam cell content of aortic sinus plaques, based on CD68 immunofluorescence (IMF; Figure 3D). After quantification, plaque composition, in terms of necrotic core and area occupied by macrophages, was indistinguishable between Tβ4^-/Y^; ApoE^-/-^ and Tβ4^+/Y^; ApoE^-/-^ (Figure 3D). These data exclude a causative difference in inflammatory response in Tβ4 mutants as being responsible for exacerbated disease progression.

**Figure 3.**
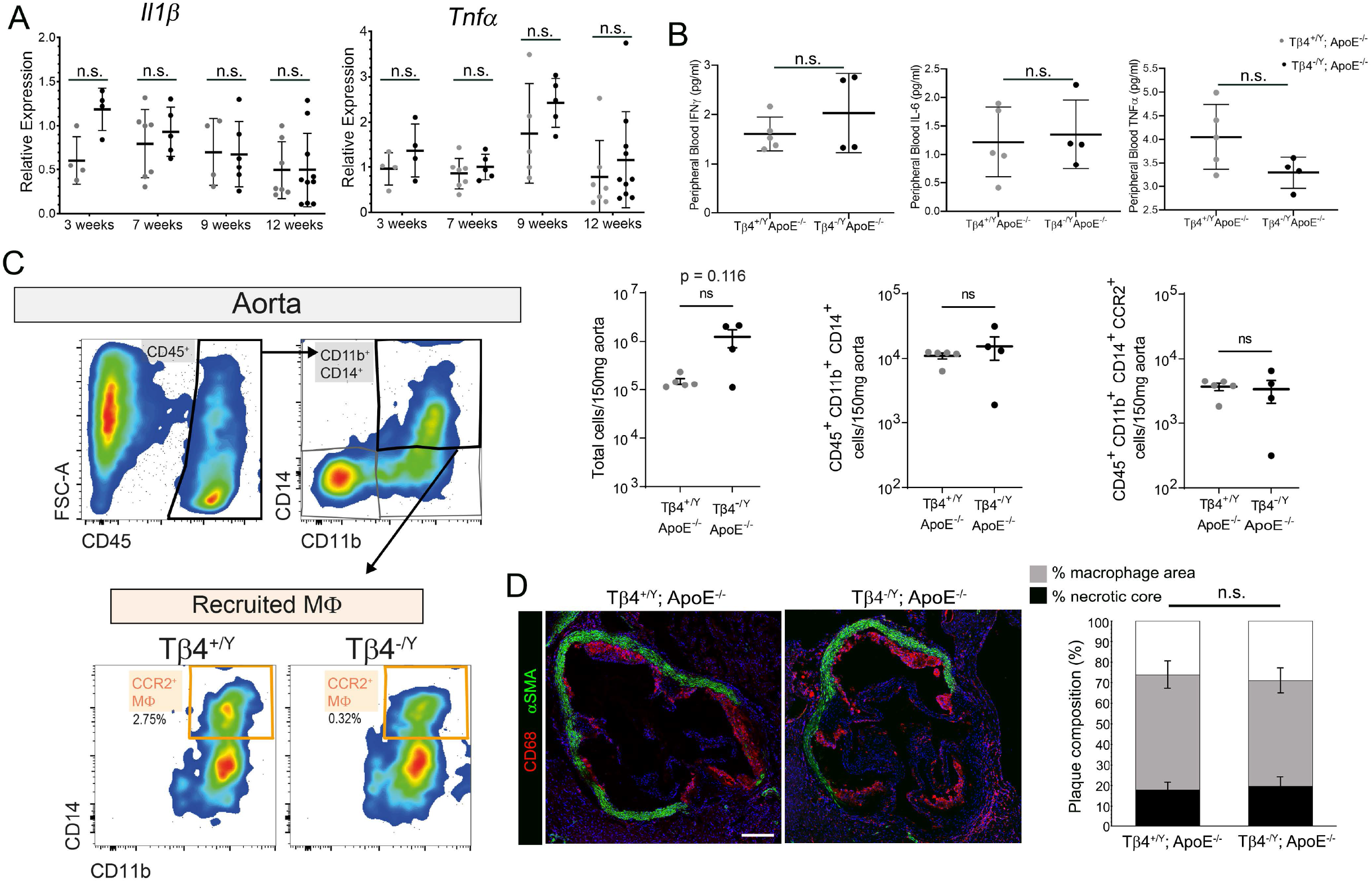
Accelerated atherosclerosis is not due to exacerbated inflammation in Tβ4-null mice. qRT-PCR of pro-inflammatory cytokines, *IIl1β* and *Tnfα*, from descending aortas of Tβ4^+/Y^; ApoE^-/-^ and Tβ4^-/Y^; ApoE^-/-^ mice (n=5 at 3 and 9 weeks’, n=6 at 7 weeks’ and n=8 at 12 weeks’ high fat diet). Serum from Tβ4^+/Y^; ApoE^-/-^ and Tβ4^-/Y^; ApoE^-/-^ mice were assayed by multiplexed automated ELISA for IFNγ, IL-6 and TNFα levels after 3 weeks’ high fat diet (n = 5; **B**). Flow cytometry to quantify macrophages (MΦ) recruited to aortas of Tβ4^+/Y^; ApoE^-/-^ and Tβ4^-/Y^; ApoE^-/-^ mice after 3 weeks of high fat diet (gating strategy shown in **C**, with the percentage of CCR2^+^ macrophages out of total live cells shown. Quantification of total cells, CD11b^+^CD14^+^ monocytes, and CCR2^+^ recruited macrophages n = 4 Tβ4^+/Y^; ApoE^-/-^ and n = 5 Tβ4^-/Y^; ApoE^-/-^ mice. Immunofluorescence of CD68+ macrophage/foam cells in aortic sinus plaques (**D**) and quantification of plaque composition, in terms of % area occupied by macrophages and % area necrotic core (n = 12). Data are presented as mean ± SD, with each data point representing an individual animal. Significance was calculated using one-way ANOVA with Tukey’s multiple comparison tests (**A**), Mann Witney non-parametric test (**B, D**), ANOVA with Bonferroni correction for multiple comparisons (**C**). n.s. = not significant. Scale bar in **D**: 250μm.

### Phenotypic modulation of VSMCs is accelerated in Tβ4-null mice

Since Tβ4 has been implicated in VSMC differentiation during development and the transcriptomic analyses above support a correlative role for the peptide in preservation of contractile phenotype, we focused our investigation on VSMC phenotype and tracked their transition as they de-differentiate in increasing numbers over the course of HFD feeding. After 7 weeks, the medial layer VSMCs of Tβ4^-/Y^; ApoE^-/-^ mice displayed a pronounced alteration in morphology, with loss of their characteristic, elongated shape and acquisition of a smaller, more rounded appearance (Figure 4A). This was associated with a shift in the ratio of synthetic over contractile markers; by IMF, a striking decrease in αSMA was accompanied by a concomitant increase in Caldesmon and Vimentin levels (Figure 4A). Tβ4^+/Y^; ApoE^-/-^ VSMCs underwent a similar shift, only this occurred more gradually, becoming noticeable from 9 weeks’ HFD, once plaques were already established (Figure 4A). By qRT-PCR, a significantly greater decrease in the ratios of contractile:synthetic genes (*Sm22α*: *caldesmon* and *Myh11*: *vimentin*) was apparent in Tβ4^-/Y^; ApoE^-/-^ mice, compared with Tβ4^+/Y^; ApoE^-/-^ (Figure 4B). Moreover, western blotting demonstrated a decrease in expression of contractile marker Calponin1 and confirmed that the overall rise in Caldesmon levels results from a proportionally larger increase in the lower molecular weight isoforms that are associated with the synthetic phenotype^6, 39^ (Figure 4C). Consistent with dedifferentiation and acquisition of a synthetic phenotype, medial VSMCs were significantly more proliferative, as determined by increased EdU incorporation between weeks 5 and 7 of HFD feeding (Figure 4D); in contrast, cells within the plaque showed no difference in proliferation between genotypes at this stage (Figure 4E). Altogether, these data demonstrate that Tβ4^-/Y^ VSMCs undergo an accelerated de-differentiation during atherosclerosis, to more rapidly become ‘synthetic’ in phenotype, compared with Tβ4^+/Y^ VSMCs, and potentially compromise vascular stability.

**Figure 4.**
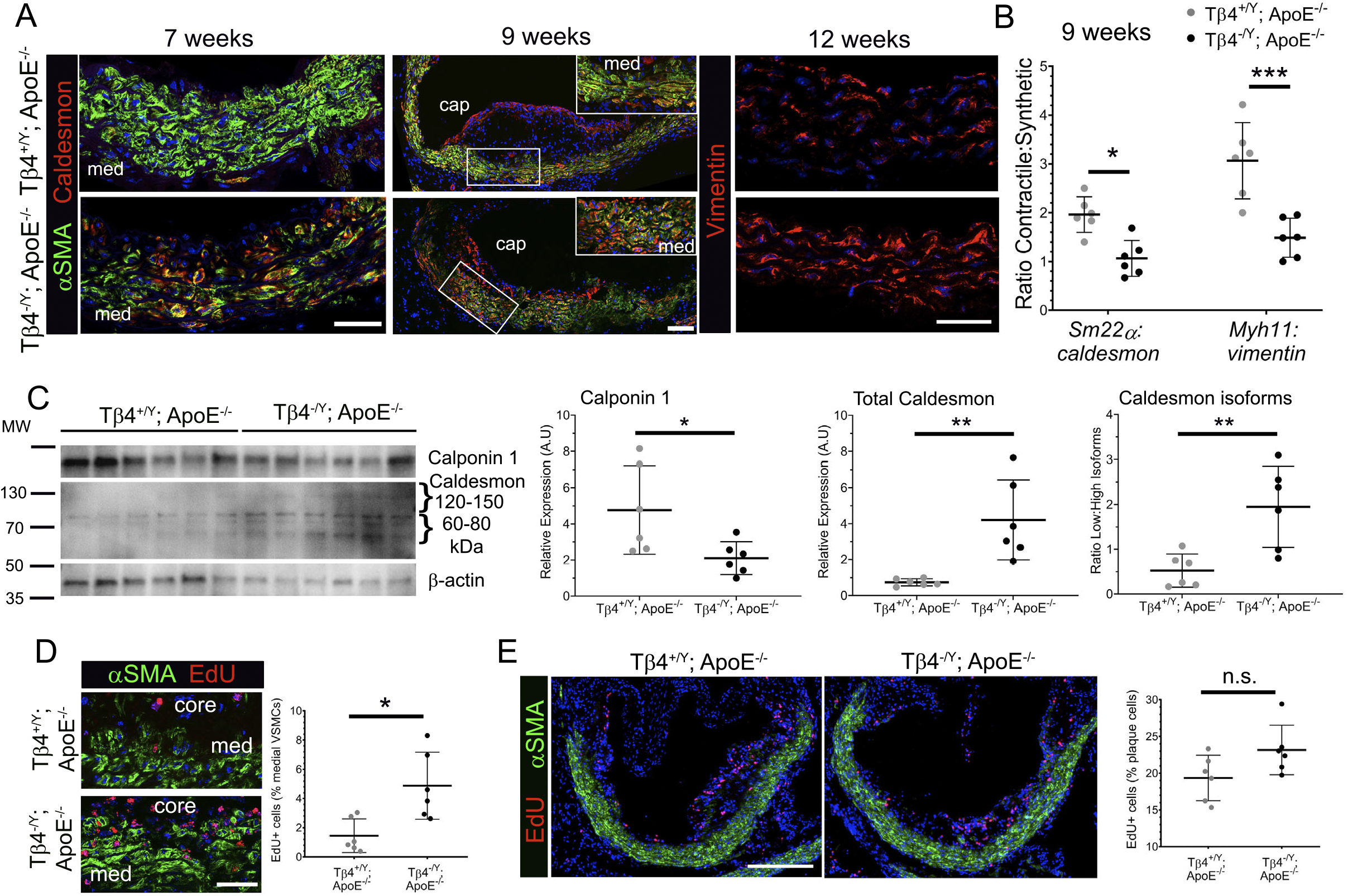
Phenotypic modulation of VSMCs is accelerated in Tβ4-null mice. Contractile-synthetic phenotype in Tβ4^+/Y^; ApoE^-/-^ and Tβ4^-/Y^; ApoE^-/-^ mice tracked over the course of lesion formation by immunofluorescence (7-12 weeks’ high fat diet, **A**). Boxed region in 9 week panel magnified in inset. Phenotype assessed quantitatively by qRT-PCR in **B** (9 weeks’ high fat diet; n = 6). Western blot showing expression of contractile proteins (high molecular weight Caldesmon, Calponin1) and synthetic markers (low molecular weight Caldesmon) in aortic lysates from Tβ4^+/Y^; ApoE^-/-^ and Tβ4^-/Y^; ApoE^-/-^ mice (**C;** n = 6), with quantification for Calponin1, total Caldesmon and low: high ratio of Caldesmon. VSMC (**D**) and plaque cell (**E**) proliferation assessed by EdU incorporation (twice weekly pulses between 5-7 weeks and harvest at 7 weeks’ high fat diet). Data are presented as mean ± SD, with each data point representing an individual animal. Significance was calculated using Mann Witney non-parametric test. n.s. = not significant; *p ≤ 0.05; **p≤0.01; ***: p≤0.001. Scale bars: **A**: 50μm (7 and 12 weeks); 100μm (9 weeks); **D**: 50μm; **E**: 250μm. med: medial layer.

### Vascular integrity and plaque stability is compromised in Tβ4-null mice

Vascular stability in the face of atherosclerotic disease is dependent upon the maintenance of viable, contractile VSMCs and the integrity of the elastin-rich matrix that they deposit. Specifically, VSMCs of the fibrous cap play an important role in ECM synthesis, to stabilize plaques and protect against rupture and thrombosis^3^. Phenotypic modulation of VSMCs frequently precedes senescence and apoptosis^40, 41^, to further destabilise affected vessels. A striking degeneration of the elastin lamellae was apparent in Tβ4^-/Y^; ApoE^-/-^ aortas, with more elastin breaks per section than in Tβ4^+/Y^; ApoE^-/-^ aortas and a higher elastin damage score (Figure 5A). By these parameters, elastin degeneration in Tβ4^-/Y^; ApoE^-/-^ aortas was comparable with, and in some cases more severe than, Myh11^Cre^; Lrp1^fl/fl^; ApoE^-/-^ aortas (Figure 5A). The serine protease high-temperature requirement factor A1(HTRA1), abundantly expressed in VSMCs^42^, was proposed to contribute towards the excessive elastolysis observed in VSMC-specific *Lrp1* KO aortas, as it depends upon LRP1-mediated endocytosis for clearance^43^. We found HTRA1 protein levels to be elevated in the medial and sub-intimal layers of Tβ4^-/Y^; ApoE^-/-^ aortas, compared with those in Tβ4^+/Y^; ApoE^-/-^ mice (Figure 5B), which may contribute towards the exacerbated elastin fragmentation and further supports the notion that endocytic clearance via LRP1 requires Tβ4. Finally, Masson’s Trichrome staining revealed extensive plaque fibrosis between 7-12 weeks (Figure 5C, shown at 12 weeks), and, while collagen content per section was greater in Tβ4^-/Y^; ApoE^-/-^ and Myh11^Cre^; Lrp1^fl/fl^; ApoE^-/-^, compared with Tβ4^+/Y^; ApoE^-/-^, it was consistently proportional to plaque size. Therefore, while mutant plaques were larger, % plaque collagen content was not disproportionately higher than control (Figure 5C). We measured fibrous cap thickness, relative to lesion thickness, after 12 weeks and, whilst this was comparable between genotypes, the caps of Tβ4^-/Y^; ApoE^-/-^ plaques were composed of cells expressing predominantly synthetic markers, such as Caldesmon (Figure 5D). In comparison, control Tβ4^+/Y^; ApoE^-/-^ cap cells retained higher levels of contractile markers, such as αSMA (Figure 5D). Of note, a significantly higher proportion of Tβ4^-/Y^; ApoE^-/-^ medial VSMCs and cells that comprise the fibrous cap were apoptotic, as revealed by TUNEL staining, compared with Tβ4^+/Y^; ApoE^-/-^ (Figure 5E). Thus, the preponderance of apoptotic and synthetic VSMCs, relative to the number of contractile VSMCs, suggests that, despite having thick fibrous caps, plaques in Tβ4^-/Y^; ApoE^-/-^ mice may be more unstable than those in Tβ4^+/Y^; ApoE^-/-^ mice. Taken together, the modulated VSMC phenotype and aggravated elastolysis are likely to destabilise the aortic wall and accelerate atherogenesis in Tβ4-LRP1 deficient mice; moreover, those plaques that form are predicted, based on lesion size as well as cap VSMC phenotype, to be more prone to rupture.

**Figure 5.**
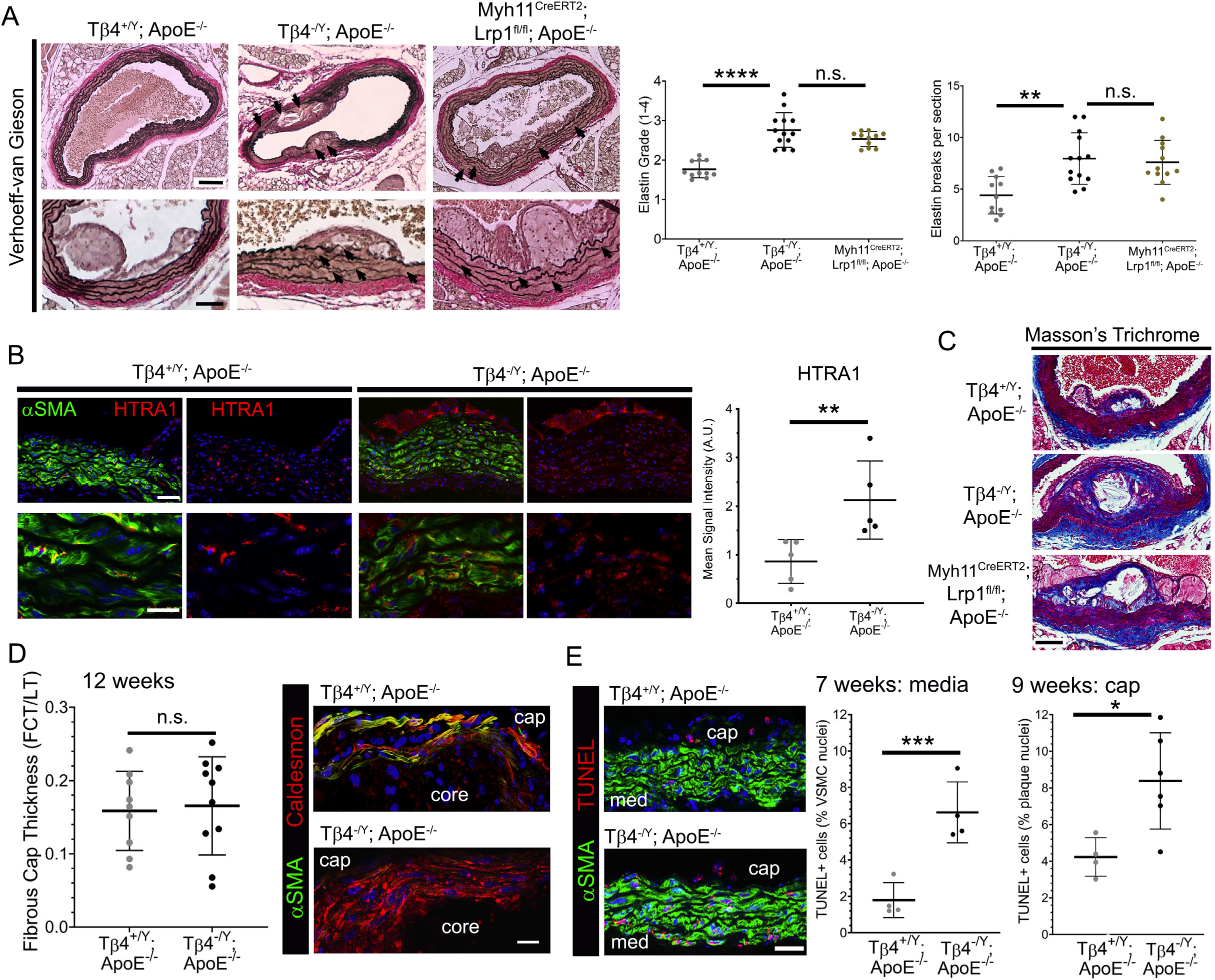
Vascular integrity and plaque stability is compromised in Tβ4-null mice. Verhoeff-van Gieson staining of abdominal aorta from Tβ4^+/Y^; ApoE^-/-^(n=11), Tβ4^-/Y^; ApoE^-/-^ (n=13) and Myh11^Cre^; Lrp1^fl/fl^; ApoE^-/-^(n=11) mice after 12 weeks’ high fat diet, to visualize elastin integrity underlying the plaque (**A**), with quantification of elastin grade (1=normal; 2=occasional breaks; 3= frequent breaks, multiple lamellae; 4=severe breakdown) and number of breaks per section. Assessment of HTRA1 accumulation in Tβ4^+/Y^; ApoE^-/-^ and Tβ4^-/Y^; ApoE^-/-^ aortic sections (n=5) after 9 weeks’ high fat diet (**B**). Fibrosis assessed by Masson’s Trichrome stain after 12 weeks’ high fat diet, representative of n=11 Tβ4^+/Y^; ApoE^-/-^, n = 13 Tβ4^-/Y^; ApoE^-/-^ and n=11 Myh11^Cre^; Lrp1^fl/fl^; ApoE^-/-^ (**C**). Fibrous cap thickness quantified, relative to lesion thickness (**D**). Assessment of apoptotic medial VSMCs and cap cells (TUNEL staining) after 7 and 9 weeks’ high fat diet (**E**). Data are presented as mean ± SD, with each data point an average of 6 sections for each animal. Significance was calculated using one-way ANOVA with Tukey’s multiple comparison tests (**A**) or Mann Witney non-parametric test (**B, D**-**E**). n.s. = not significant; *p ≤ 0.05; **p≤0.01; ***: p≤0.001; ****p≤ 0.0001. med: medial layer; FCT: fibrous cap thickness; LT: lesion thickness. Scale bars: **A**: 200μm (upper); 100μm (lower); **B**: 50μm (upper) 10μm (lower); **C**: 100μm; **D**: 20μm; **E**: 50μm.

### LRP1-controlled growth factor signaling is dysregulated in Tβ4-null mice

The protective, VSMC-autonomous function of LRP1 has been attributed to the endocytic control of PDGFRβ signalling, as well as to the endocytic clearance of matricellular proteins that are responsible for ECM remodelling^43, 44^. Our demonstration that Tβ4^-/Y^ phenocopy VSMC-specific LRP1 KO, in their susceptibility to atherosclerotic plaque formation, aggravated elastolysis and compromised clearance of HTRA1, raises the possibility that defects in Tβ4 mutants are, similarly, due to LRP1-PDGFRβ pathway dysregulation. We therefore assessed readouts of the LRP1-PDGFRβ pathway in medial layer VSMCs at early-intermediate disease stages, which coincide with initiation of injury-provoked phenotypic switching. Tyr1021 PDGFRβ phosphorylation, which reports the extent of receptor autophosphorylation following ligand binding and LRP1-mediated endocytosis^45^, was elevated 2.4-fold in VSMCs of Tβ4^-/Y^; ApoE^-/-^ aortas, compared with Tβ4^+/Y^; ApoE^-/-^ aortas, by immunofluorescence (Figure 6A). Similarly, levels of Tyr 4507 phospho-LRP1, resulting from phosphorylation by PDGFRβ^46,47^, were elevated 1.8-fold (Figure 6A). Receptor phosphorylation is coupled via adaptor protein binding and phosphatidylinositol 3-kinase (PI3K) activation to downstream effectors, including AKT and p42/p44 MAP kinase (ERK1/2)^48^. By western blotting, phosphorylated (activated) AKT and p42/p44 MAP kinase were significantly higher in aortas of Tβ4^-/Y^; ApoE^-/-^ aortas, compared with Tβ4^+/Y^; ApoE^-/-^ aortas, by immunofluorescence (Figure 6B). These data indicate that accelerated phenotypic modulation of VSMCs and exacerbated atherosclerosis in Tβ4^-/Y^ mice is associated with dysregulated LRP1-PDGFRβ pathway activity and point to a likely role for Tβ4 in controlling receptor turnover, as previously elucidated in isolated VSMCs^15^, to influence the progression and outcome of atherosclerotic disease.

**Figure 6.**
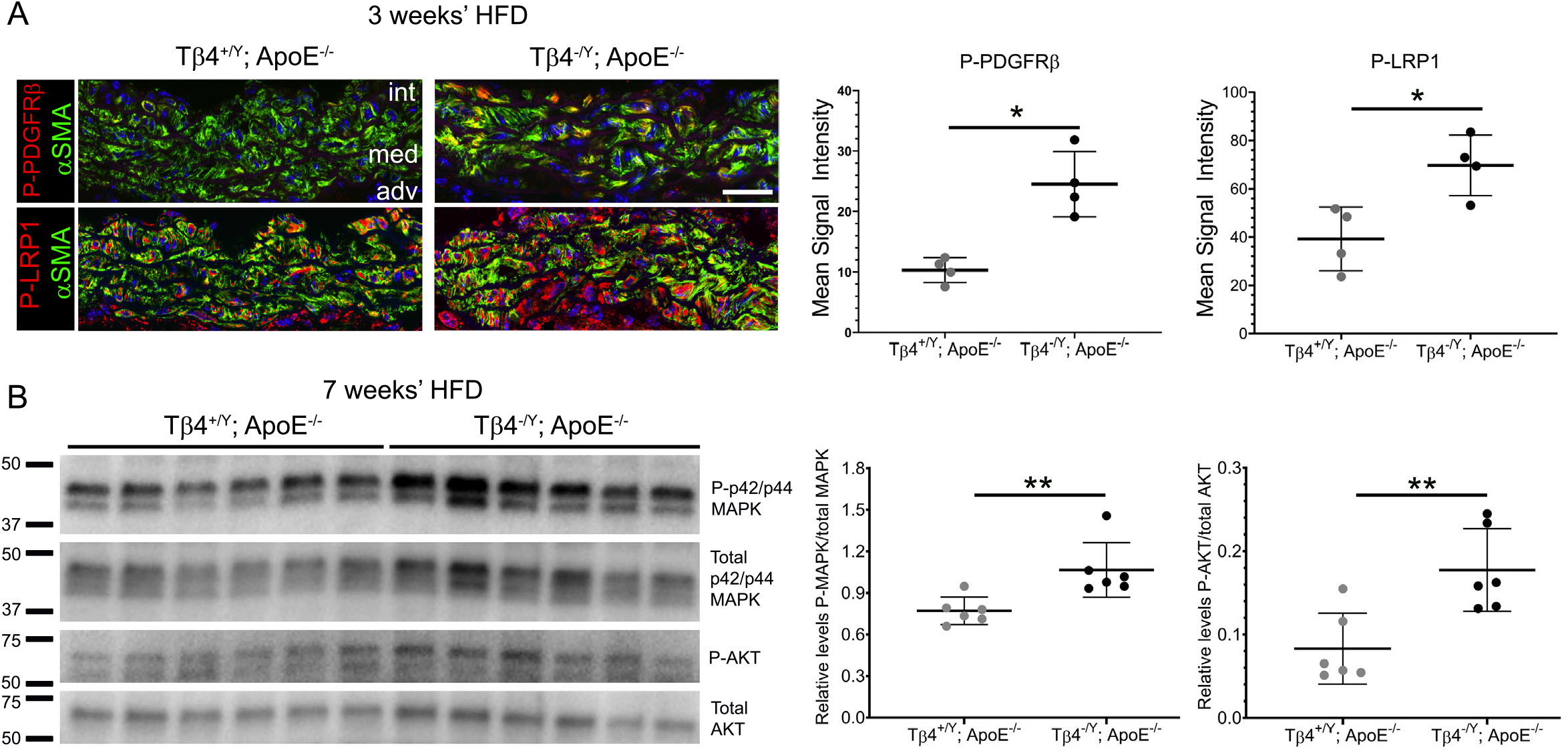
LRP1-controlled growth factor signaling is dysregulated in Tβ4-null mice. Phospho-LRP1 (Tyr 4507) and Phospho-PDGFRβ (Tyr1021) levels in medial VSMCs of Tβ4^+/Y^; ApoE^-/-^ and Tβ4^-/Y^; ApoE^-/-^ (n=4) after 3 weeks’ high fat diet (**A**). Western blotting of downstream effectors, phosphorylated p42/p44 MAPK and AKT, expressed relative to total p42/p44 MAPK and AKT from n=6 Tβ4^+/Y^; ApoE^-/-^ and Tβ4^-/Y^; ApoE^-/-^ aortas after 7 weeks’ high fat diet (n = 6; **B**). Data are presented as mean ± SD, with each data point representing an individual animal. Significance was calculated using a Mann Witney non-parametric test. * P≤ 0.05; ** P ≤ 0.01. int: intima; med: media; adv; adventitia. Scale bars: **A**: 50µm.

## Discussion

Collectively, our study defines a key role for Tβ4 to maintain medial VSMCs in a differentiated, contractile state to protect against atherosclerosis. Tβ4 null mice on an atherogenic (ApoE^-/-^) background develop large complex plaques, irrespective of gender, indicating that loss of Tβ4 exacerbates disease progression. Assessment in both genders was important, due to the increasing reports of sexual dimorphism, both in patients and in experimental studies of cardiovascular disease^34^. Accelerated disease progression was characterized by augmented VSMC phenotypic modulation and underpinned by dysregulated PDGFRβ signaling and deleterious ECM remodelling, both of which may result from a failure of Tβ4 to functionally regulate LRP1 trafficking. Mechanistically, Tβ4 has been shown to bind the intracellular domain of LRP1, near NPxY domains that are necessary for signal transduction and endocytosis; accordingly, Tβ4 controls the balance between recycling and lysosomal targeting of LRP1-PDGFRβ complexes, to critically regulate VSMC sensitivity and responses to PDGF-BB^15^. Consistent with this, vascular defects in Tβ4 null mice closely phenocopy those reported with VSMC-specific loss of LRP1^13, 20, 33^, not only at baseline and in AAA^15^, but also, herein, in atherosclerotic disease. Thus, our findings further add support to the existence of a common regulatory mechanism through which Tβ4 and LRP1 act in vascular homeostasis to maintain vessel stability and protect against disease.

The significance of VSMC phenotypic switching in atherosclerosis is still not fully resolved^3^. While abnormal proliferation and migration of VSMCs drives early lesion formation and, debatably, may contribute pro-inflammatory macrophage-like cells to the plaque^23, 49, 50^, VSMC-derived ‘fibromyocytes’ are beneficial in stabilising more advanced plaques to prevent fibrous cap rupture. Thereafter, however, prolonged VSMC transformation perpetuates the inflammatory environment and promotes pathogenic remodelling, causing intimal thickening, arterial stiffness and stenosis^9^. Understanding VSMC heterogeneity, and identifying the regulators of phenotypic modulation may enable the fine tuning of VSMC phenotypes that are beneficial for repair. Transcriptomic and proteomic analyses at single cell resolution make this an attainable goal within the next decade.

A limitation of this study is that global deletion does not allow us to distinguish cell type-specific roles for Tβ4. While it was important to exclude inflammation as the primary reason for accelerated disease progression in Tβ4^-/Y^; ApoE^-/-^ mice, given the array of anti-inflammatory roles ascribed to Tβ4^35, 37, 51^ and LRP1^52^, a comprehensive immune cell profiling is required to categorically exclude roles beyond macrophage recruitment. Although correlative, the scRNA-seq analysis and validation in plaque sections strongly supports a cell-autonomous requirement for Tβ4 in VSMCs for preservation of differentiated contractile phenotype. When taken together with the prior demonstration of baseline defects and AAA predisposition after VSMC-specific depletion of Tβ4^15^, an endogenous role within VSMCs appears, overall, most likely, but still requires further investigation.

LRP1 is increasingly recognised for its central role in protection against cardiovascular disease, with mutations in the gene now implicated through genome-wide association studies as a causal factor in AAA^17^, carotid artery^18^ and coronary artery disease^19^. From these studies, it appeared logical that the genetic predisposition should result from defects, either in the removal of proatherogenic lipids via the liver or in exacerbated LDL uptake by macrophage and conversion to foam cells^53^. However, animal studies were instrumental in enabling selective targeting in macrophages^54^, endothelial cells^55^, VSMCs^56^, perivascular adipose tissue^57^ and hepatocytes^58^ to discriminate roles for LRP1 in the respective cell types. While LRP1 is instrumental in macrophages to maintain cholesterol efflux^59^ and enable efferocytosis^60, 61^, distinct, vasculoprotective functions were defined for LRP1 in VSMCs, predominantly in the control of growth factor signalling and ECM remodelling^13, 20, 56^. That loss of Tβ4, a molecular regulator of LRP1 endocytosis, recapitulates the phenotype of LRP1-depleted VSMCs emphasises the importance of controlled receptor trafficking in determining medial VSMC responses and progression of AAA and atherosclerosis.

## Author contributions

SM, ANR and NS carried out most experiments and data analysis, with computational analysis of scRNA-Seq data by ANR and BP. NS established the hypotheses, supervised the study and wrote the manuscript. All authors edited and approved the manuscript.

## Acknowledgements

The study was funded primarily by the British Heart Foundation Ian Fleming Senior Basic Science Research Fellowship (FS/13/4/30045), awarded to NS, and also by a studentship from the Oxford Medical Research Council Doctoral Training Partnership to SM.

## Figure Legends

**Online Figure I. scRNA-seq of VSMCs in atherosclerosis**.

**A**: Computational analysis of scRNA-seq of lineage traced aortic VSMCs from ApoE^-/-^ mice (derived from^23^) identified 5 VSMC subpopulations, as shown in Figure 1. Heatmap illustrating the top 5 marker genes and their expression levels to characterise each cluster. **B**: Stacked bar plot depicting the proportions of each VSMC subpopulation at each time point, to illustrate VSMC modulation in atherosclerosis. **C**: Violin plots to demonstrate changes in *Tmsb4x* and *Lumican* expression across the five VSMC subpopulations over the disease time course.

**Online Figure II. Tβ4**^**+/Y**^**; ApoE**^**-/-**^ **and Tβ4**^**-/Y**^**; ApoE**^**-/-**^ **do not show differences in weight gain or cholesterol levels after western diet**. Comparison of weight gain (**A**) and plasma cholesterol levels (**B**) in Tβ4^+/Y^; ApoE^-/-^ and Tβ4^-/Y^; ApoE^-/-^ mice over the time course of the high fat diet feeding regime, compared with Myh11^Cre^; Lrp1^fl/fl^; ApoE^-/-^ mice. Data are presented as mean ± SD, with n=9-11 in **A**. Each data point in **B** represents an individual animal. Significance was calculated using two-way ANOVA with Bonferroni correction for multiple comparisons (**A**) and one-Way ANOVA with Tukey’s multiple comparison tests (**B**). n.s. = not significant; ***: p≤0.001

**Online Figure III. Increased predisposition to atherosclerotic plaque formation was also observed in female Tβ4**^**-/-**^**; ApoE**^**-/-**^ **mice**. *En face* aorta preparations and oil red O staining to visualize plaques in female mice fed Western diet for 12 weeks (**A**). Quantification of plaque coverage in the aortic arch, thoracic aorta (TA), abdominal aorta (AA) and total aorta (**B**). Weight gain was significantly increased in female Tβ4^-/-^; ApoE^-/-^, compared with Tβ4^-/-^; ApoE^-/-^ mice from 4-12 weeks’ high fat diet feeding (**C**). Data are presented as mean ± SD, with each data point representing an individual animal (**B**) and n=8 (**C**). Significance was calculated using one-way ANOVA with Tukey’s multiple comparison tests (**B**) and two-way ANOVA with Dunnett’s post hoc tests (**C**). n.s.= not significant; *p ≤ 0.05; **p≤0.01; ***: p≤0.001.

