## Supplementary figures and images for "Thymosin β4 preserves vascular smooth muscle phenotype in atherosclerosis via regulation of Low Density Lipoprotein Related Protein 1 (LRP1)"

### Supplemental figures

Online Figure I

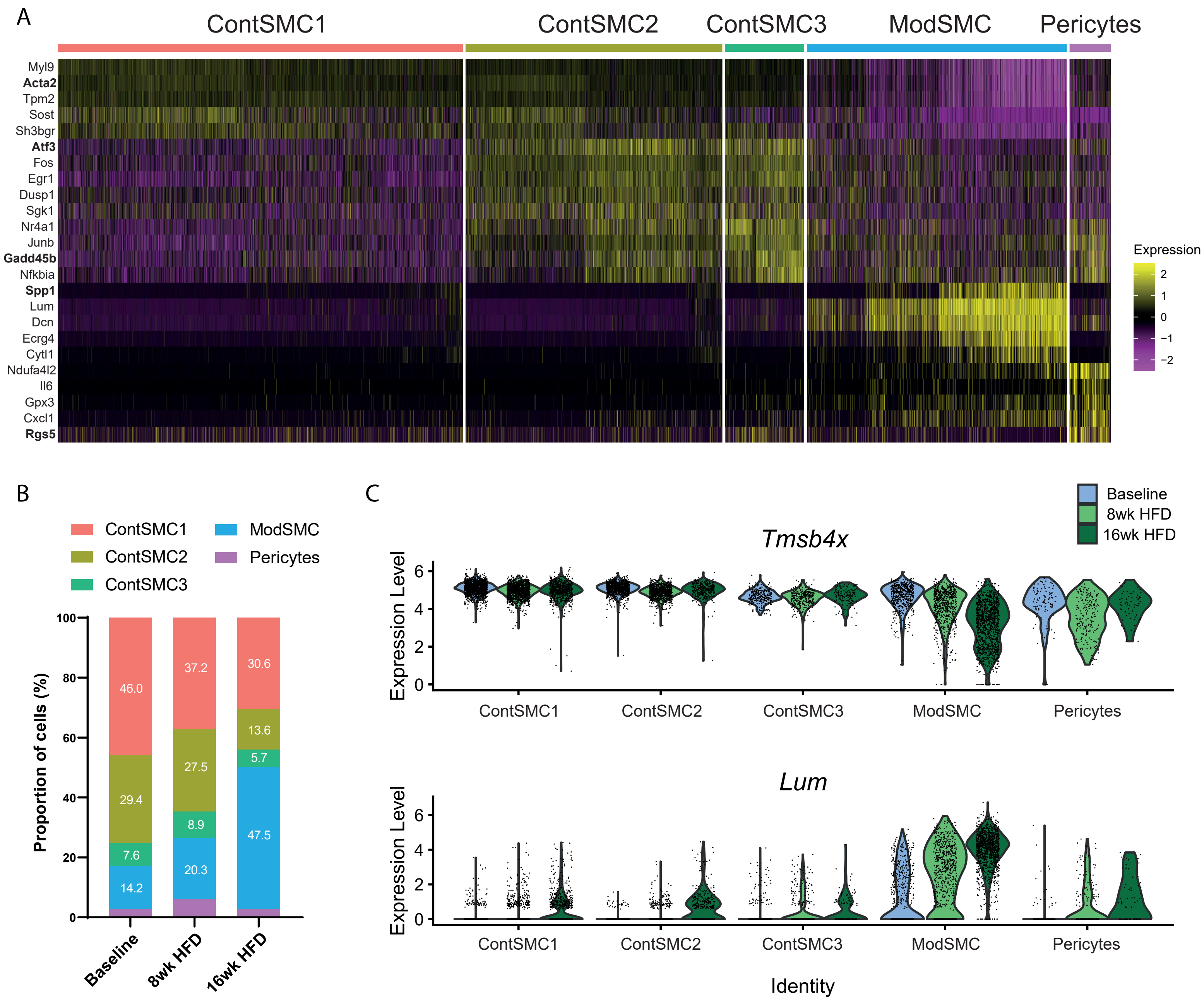

Online Figure II

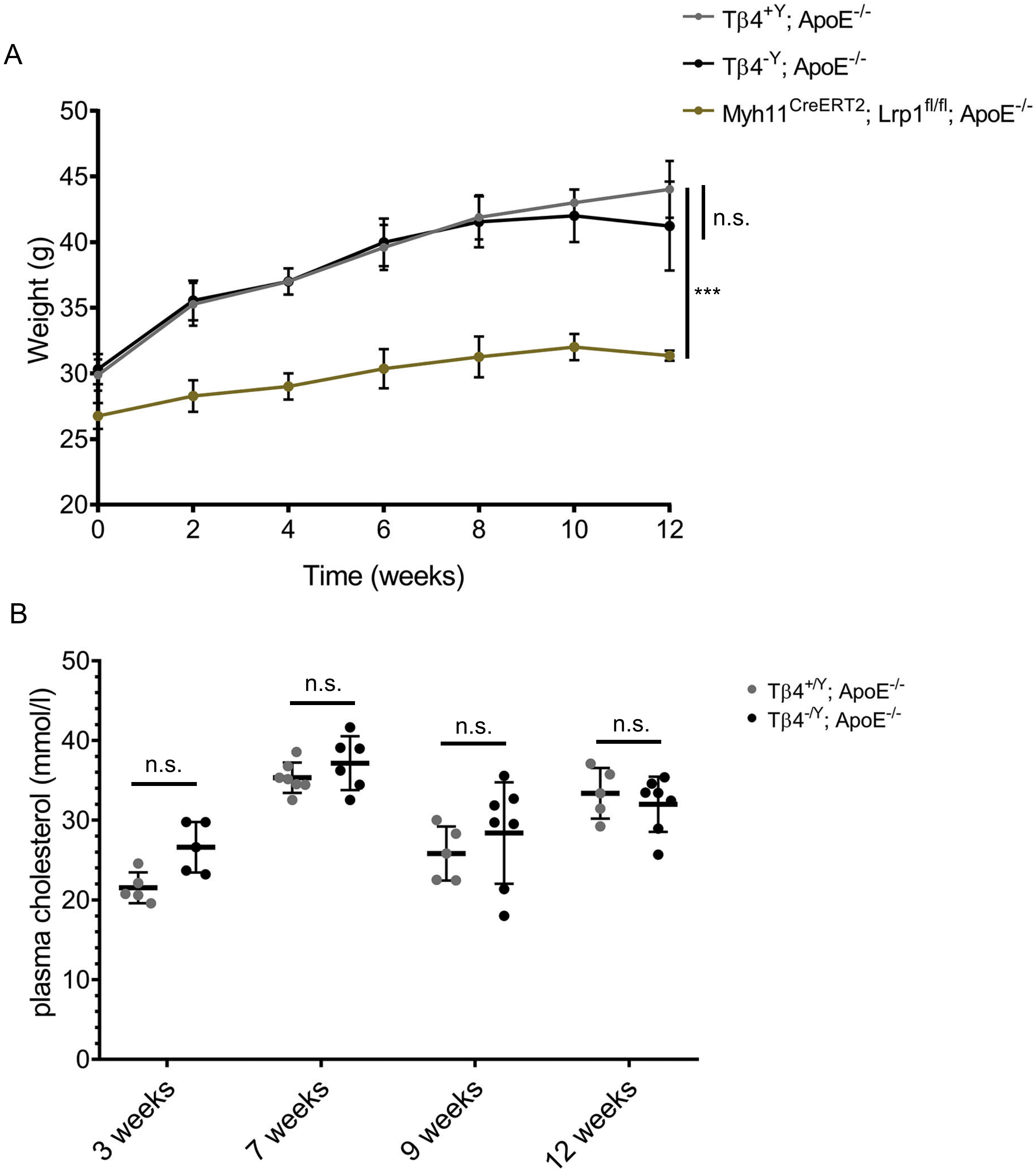

Online Figure III

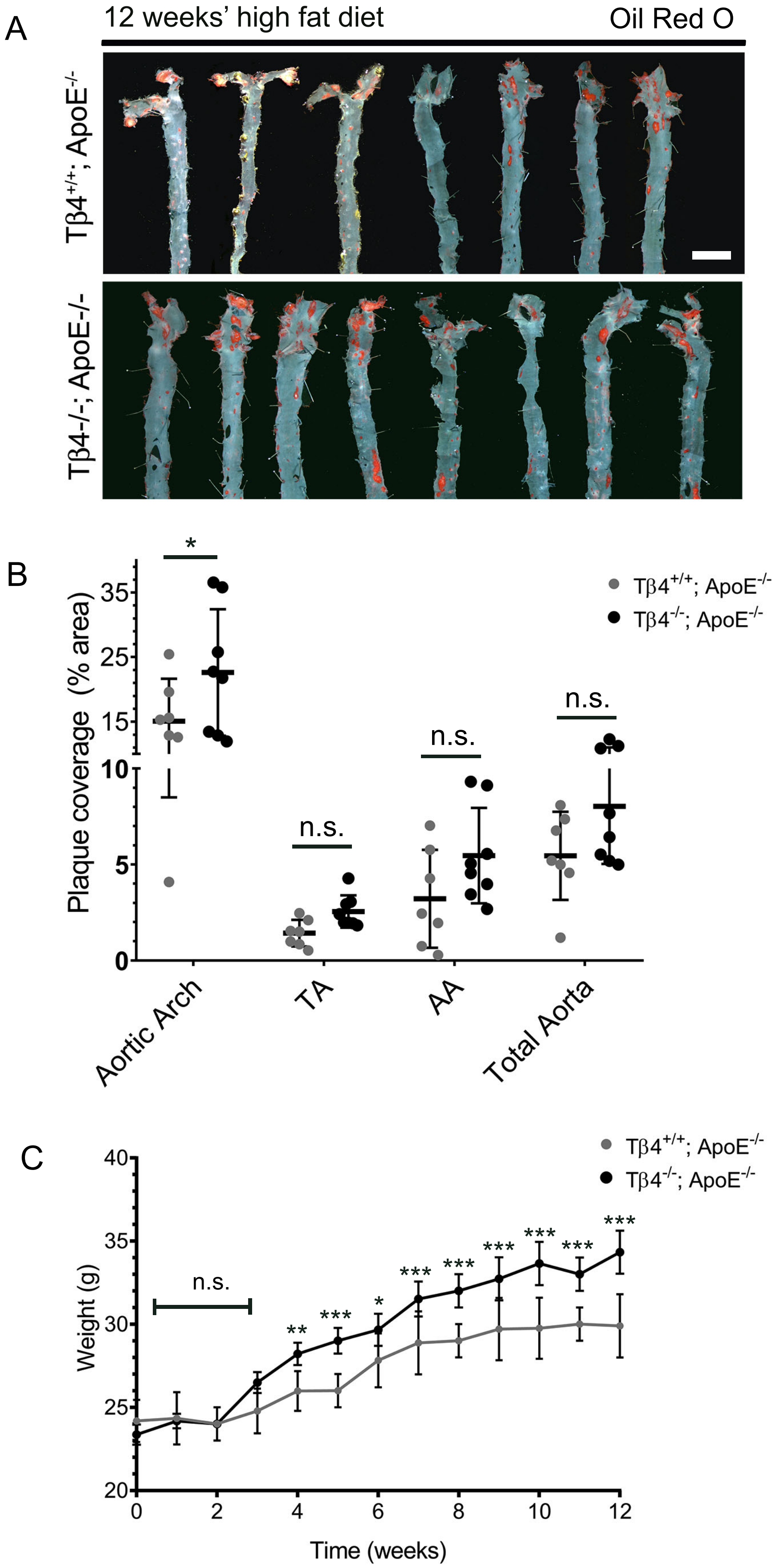
